# Seasonal changes in recombination rate, crossover interference, and their response to desiccation stress in a natural population of *Drosophila melanogaster* from India

**DOI:** 10.1101/2020.06.17.156877

**Authors:** Dau Dayal Aggarwal, Sviatoslav Rybnikov, Shaul Sapielkin, Eugenia Rashkovetsky, Zeev Frenkel, Manvender Singh, Pawel Michalak, Abraham B. Korol

**Affiliations:** Department of Zoology, Banaras Hindu University, Varanasi 221005, India; Institute of Evolution, University of Haifa, 3498838 Haifa, Israel; Department of Evolutionary and Environmental Biology, University of Haifa, 3498838 Haifa, Israel; Department of Biotechnology, UIET, MD University, Rohtak 124001, India; Edward Via College of Osteopathic Medicine, Blacksburg, VA 24060, USA; Center for One Health Research, Virginia-Maryland College of Veterinary Medicine, Blacksburg, VA 24060, USA

**Keywords:** recombination rate, crossover interference, indirect selection for recombination, seasonality, plasticity, desiccation, *Drosophila melanogaster*

## Abstract

Environmental seasonality is a potent evolutionary force, capable to maintain polymorphism, promote phenotypic plasticity, and cause bet-hedging. In *Drosophila*, it has been reported to affect life-history traits, tolerance to abiotic stressors, and immunity. Oscillations in frequencies of alleles underlying fitness-related traits were also documented alongside SNP alleles across genome. Here, we test for seasonal changes in recombination in a natural *D. melanogaster* population from India using morphological markers of the three major chromosomes. We show that winter flies (collected after the dry season) have significantly higher desiccation tolerance than their autumn counterparts. This difference proved to hold also for hybrids with three independent marker stocks, suggesting its genetic rather than plastic nature. Significant segment-specific changes are documented for recombination rate (in five of 13 intervals) and crossover interference (in five of 16 studied pairs of intervals); both single- and double-crossover rates tended to increase in the winter cohort. The winter flies also display weaker plasticity of recombination characteristics to desiccation. We ascribe the observed differences to indirect selection on recombination caused by directional selection on desiccation tolerance. Our findings suggest that changes in recombination can arise even after a short period of seasonal adaptation (~8–10 generations).

## 1. Introduction

Environmental seasonality plays an important role as an ecological factor, and its significance as a potent evolutionary force is becoming increasingly evident [1]. The evolutionary consequences of within-year oscillations in selection directions and intensities considerably depend on the generation time of the species in question. In perennials, exposure to environmental seasonality as lifespan-long regular background may select for pleiotropy and phenotypic plasticity. In annuals, whose developmental stages are distributed across the year, it may additionally select for time-tuning of life-history traits and bet-hedging. Yet, the most interesting organisms from the evolutionary viewpoint are multivoltine species, having several generations per year. Here, seasonality may lead, in addition to all the above-mentioned adaptations, to far-reaching population-level effects, including maintenance of balanced polymorphism [2–4], complex dynamics of allele frequencies [5,6], and evolving dominance [7,8]. Moreover, multivoltine species seem to be the most appropriate models for addressing the intriguing interplay between different adaptations to seasonality, including the interaction between plastic and heritable responses to periodical environmental stressors.

Fruit flies are one of the most appropriate models for seasonality studies. The population size of various *Drosophila* species has long been known to fluctuate during the year [9,10]. Later studies have also revealed seasonal oscillation in several important fitness-related phenotypic traits, including desiccation tolerance [11–13], the activity of metabolic enzymes [14], life-history traits, resistance to heat, cold and starvation [15], and innate immunity [16]. In their recent extensive genome-wide analysis, Bergland *et al*. [17] have revealed hundreds of seasonally fluctuating SNPs; the authors relate them to variation in adaptive phenotypic traits, first of all cold- and starvation tolerance.

Meiotic recombination is a key source of genetic variation in sexually reproducing organisms; its variation may have important evolutionary consequences, including for *Drosophila* [18,19]. If seasonal changes in the above-mentioned fitness-related traits are, at least partially, a result of genetic adaptation, a natural question is whether selection on these traits can give rise to temporal indirect selection on recombination. We address this question in the current study, by comparing the recombination characteristics of two seasonal cohorts obtained from the same natural population of *D. melanogaster*. We hypothesize that if the studied population has evolved a higher desiccation tolerance during the dry winter, this will cause accompanying changes in recombination characteristics. The possibility of indirect selection for recombination has been proven in several experimental-evolution studies, including our own [20], where a segment-specific increase in both single- and double-crossover rates was observed after 48 generations of directional selection for desiccation tolerance. Moreover, similar changes have been observed after much shorter periods of selection for recombination-unrelated traits, e.g., after 15 generations in fruit fly [21] and just five generations in cabbage [22]. However, changes in recombination during seasonal adaptation of a natural population have never been studied before, to the best of our knowledge.

We consider two recombination characteristics: recombination rate (RR) and crossover interference (CI), the ability of one crossover event to affect the probability of crossover events in other regions. We also examine the plasticity of both characteristics in response to desiccation stress and test whether the plasticity differs between the seasonal cohorts. Since the pioneering works by Plough [23,24], both RR and CI have been recognized as traits exhibiting considerable plasticity to environmental stressors; see also [25–27]. Our recent study [28] has shown that in terms of both recombination characteristics (RR and CI), desiccation-tolerant lines from the above-mentioned experiment [20]) respond to desiccation stress weaker than desiccation-sensitive lines. Here, we investigate, for the first time, seasonal variation in the plasticity of RR and CI in a natural population.

Importantly, recombination characteristics do not directly affect the individual’s survival or reproductive success, in contrast to stress tolerance or other fitness-related traits. However, they do affect important population-level features, such as mean fitness, genetic variation, genetic load and speed of adaptation, which suggests that recombination variation can be adaptive [29–31]. Several studies have found that recombination may vary in natural populations along certain spatial environmental gradients [32–34]. Such findings argue, despite their scarcity, for the adaptivity of recombination variation. The seasonal changes in recombination analysed in this study represent another, *temporal* aspect of recombination variation. To date, such changes were addressed during either an individual’s lifespan [35–37] or a population’s phase-transition [38], but not during a population’s adaptation to its seasonal environment.

## 2. Material and Methods

### (a) Flies and crosses

The flies were collected using the net-sweeping method in the wild, at the lowland locality Kalka in Western Himalaya, India. The samples were collected at the end of climatic autumn (September) and at the end of climatic winter (February). The two seasons significantly differ in temperature and relative humidity: the autumn is warm and wet while the winter is cool and dry [39]. For each season, we established seven isofemale lines. Before the recombination-assessment experiment, the flies were maintained during five to six generations on the yeast-agar-sugar food medium at temperature 22°C and relative humidity 65-70% (hereafter “normal conditions”).

For the recombination-assessment experiment, 20 virgin females (6-day-old post eclosion) from each isofemale line were allowed to mate with males from three marker stocks, containing visually distinguishable morphological markers in one of the large chromosomes. In total, we examined 16 markers: four in chromosome X (*y, cv, v*, and *f*), seven in chromosome 2 (*al, dp, b, pr, c, px*, and *sp*), and five in chromosome 3 (*ru, h, th, sr*, and *e*).

RR and CI were assessed in the obtained F_1_-hybrids, heterozygous for the studied markers. Hereafter, they are referred to as “the autumn hybrids” and “the winter hybrids’, depending on the origin of their parental lines. In each of 42 crosses (seven autumn parental lines and seven winter parental lines, each crossed with three independent marker stocks), virgin females were divided into the treatment and the control groups, each of 20-22 individuals. The treatment group was subjected to desiccation hardening, short-term sub-lethal desiccation stress (see Subsection 2b), while the control group was kept growing in the normal conditions. Upon the treatment (if imposed), the females were transferred to fresh food medium and allowed to mate with males from the corresponding marker stock for the next four days. After mating, the females were allowed to lay eggs in fresh bottles for 48h. The obtained test-cross progeny was scored for the morphological markers to assess the RR and CI.

### (b) Desiccation-tolerance estimation and desiccation hardening

Desiccation tolerance was quantified as the time till lethal dehydration of all flies in dry air (LT_100_). Ten virgin females (six days after eclosion) were placed into a dry plastic vial containing 2g of silica gel at the bottom. The vials were covered with a disk of foam and then placed into a desiccator chamber (Secador electronic desiccator cabinet; www.tarson.com) maintaining the relative humidity of 5–8%. The number of immobile flies was scored every 30 min during the first ten hours, and every 15 minutes thereafter. We used ten replicates per isofemale line, resulting in 70 replicates per season.

Flies in the treatment group were subjected to desiccation hardening, a short-term exposure to dry air resulting in 5% mortality. The value of LT_5_ was calculated using the probit analysis, separately for each cross.

### (c) Statistical analysis

The experiments include 14 lines, seven collected in autumn season (**a**) and seven collected in winter season (**w**). Each wild-type line was crossed to lines marked for chromosomes X, 2, and 3; the resulted F_1_-females were either subjected to desiccation hardening (treatment, **t**) or reared in normal conditions (control, **c**). Using the segregating test-cross progeny, we were interested to test the effects of season, treatment, line, and their interactions on RR and CI. The analysis was based on maximum-likelihood (ML) approach, in the form of: (*i*) logistic regression with logit link function for testing the effects of the mentioned factors on RR; (*ii*) restricted ML analysis for the effects on CI.

### Variation in recombination rate

Analysis of the effect of season on RR was performed separately for each marker interval. For comparison of the progeny from non-treated females we employ generalized linear model (GZLM in *Statistica* software) for binomial data (crossovers vs non-crossovers in the testcross progeny) using logit link function. In this analysis, we tested for the effects of ‘seasons’ and ‘lines’ nested in seasons in accordance to the following approximation:

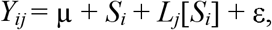

where Y*_ij_* represents the logit-transformed observed proportion of crossovers in the considered interval in line *Lj* (*j*=1,…,7) from season *S_i_*, i.e. autumn (a) or winter (w), and μ is the general mean. Similarly, the effect of treatment on RR, separately for each of the two seasons, was analysed using GZLM with the representation:

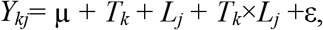

where *Y_kj_* is the logit-transformed observed proportion of crossovers in the considered interval in line *j*=1,…,7 from treatment *T_k_*, i.e. control (c) or treatment (t), and μ is the general mean.

### Variation in crossover interference

This part of data treatment and inference was based on direct ML analysis, similarly to [20,28]. In the analysis of CI, we employed an approach that can be referred to as restricted ML, since we use here the estimates of RR calculated for single intervals for each of the 28 season-line-treatment combinations separately. Considering these estimates as constant parameters rather than variables simplifies inference of the season, line and treatment effects on CI (coefficient of coincidence, *C*). Thus, our ML analyses include four vectors of parameters to be estimated and compared:

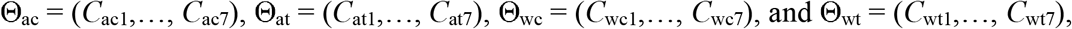

where the indices ac, at, wc and wt stand for *autumn-control*, *autumn-treatment*, *winter-control,* and *winter-treatment* combinations, respectively. In general, for a pair of consequent intervals flanked by a trio of markers (m1-m2-m3), the log-likelihood function for a sample from a testcross progeny can be presented as

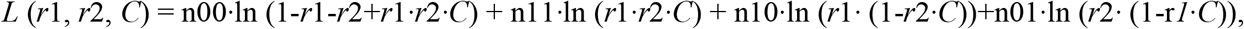

where *r*1, *r*2, and *C* are the (unknown) crossover rates in m1-m2 and m2-m3 intervals and *C* is the coefficient of coincidence, while n00 is the observed number of non-recombinants for both intervals, and n10, n01 and n11 are the numbers of recombinants for only the first, only for the second, and for both intervals, respectively. In our estimation of parameter *C,* we maximize the likelihood function under restriction that the values of *r*1 and *r*2 are already calculated in single interval analysis [40]. We do it avoid situations when the estimate of RR for a certain interval obtain in 3-locus analysis depends on the second interval.

*<u>Season effects for non-treatment data</u>*: in this situation, the restricted log-likelihood functions depend on vectors of seven variables, Θac = (*C*_ac1_,…, *C*_ac7_) and Θ_wc_ = (*C*_wc1_,…, *C*_wc7_). Our H0 hypothesis is that CI does not depend neither on seasons nor on lines within seasons, i.e. *C*_ac1_=…=*C*_ac7_ = =*C*_wc1_=…= *C*_wc7_. Our H1 hypothesis is that C depends only on season, i.e. *C*_ac1_=…=*C*_ca7_, and *C*_wc1_=…= =*C*_wc7_. Our H2 hypothesis is that C depends both on season and on lines, i.e. in general case we need all 14 parameters. ML estimates for H0, H1, and H2 include optimization for 1, 2, and 14 parameters respectively. To discriminate between these hypotheses, we use likelihood-ratio (LR) test with df=1, 12 and 13 for pairs H0-H1, H1-H2, and H0-H2, respectively.

*<u>Stress-treatment effect within season:</u>* given the season and line effects on CI, we tested if treatment had an effect. Here our H0 hypothesis assumes no difference in CI between control and treatment for each of the seven tested lines within a season (implying seven parameters in the model per season). Hypothesis H1 is that CI changes under treatment (implying 14 parameters in the model per season). The standard LR test with df=14-7=7 enables to discriminate between H0 and H1. The standard test is not sensitive to the direction of induced changes in *C*. As a result, if changes in all lines are high but oppositely directed, then H0 will be rejected even in the absence of an overall directed treatment effect. To overcome this problem, the following test was used.

Let Y= Σ Y_line_/sqrt(7), where Y_line_=sqrt(X_line_^2^)*sign(C_line_treatment -C_line_control) and X_line_^2^ = 2(log*L*(H1)-log*L*(H0)). Under H0, X_line_^2^ has χ2 distribution with df=1. In case of no consistent direction of treatment effect (i.e. the treatment-control differences across lines within season have symmetric distribution), Y_line_ value has asymptotically normal distribution. Hence, under H0 and the mentioned symmetry, the statistic Y is also normally distributed. If the absolute value of Y is lower than the critical value even though X_total_^2^ was higher than the critical value, we conclude that the significance of heterogeneity of stress response of *C* values was caused by heterogeneity of lines’ response direction rather than the overall direction of response.

## 3. Results

### (a) Desiccation tolerance

We found that the winter parental lines had, on average, higher desiccation tolerance (measured as LT_100_) than the autumn ones: 28.31±1.30h against 22.59±0.88h (Fig. 1). The difference in means was highly significant: Mann-Whitney *U*=46.0, *p*=0.004. Remarkably, the same pattern held for hybrids with each of the three marker stocks: the winter hybrids had significantly higher desiccation tolerance than the autumn ones (*U*≥47.0, *p*≤0.002). The difference between the hybrids manifested also in terms of LT_5_: it was, on average, 4h 15min and 4h 30min for the autumn and the winter hybrids, respectively. With this respect, the latter were subjected to 15min longer desiccation hardening in the recombination-plasticity assay. Notably, four different estimates of desiccation tolerance (measured in the parental lines and their hybrids with the three marker stocks) appeared highly concordant: Spearman’s rank correlation *ρ*≥0.84, *p*≤2·10^−4^ (Fig. 1). Besides, for all these estimates, the studied seasonal cohorts had the same variance: non-parametric Levene’s criterion *F*≤0.079, *p*≥0.784).

**Figure 1.**
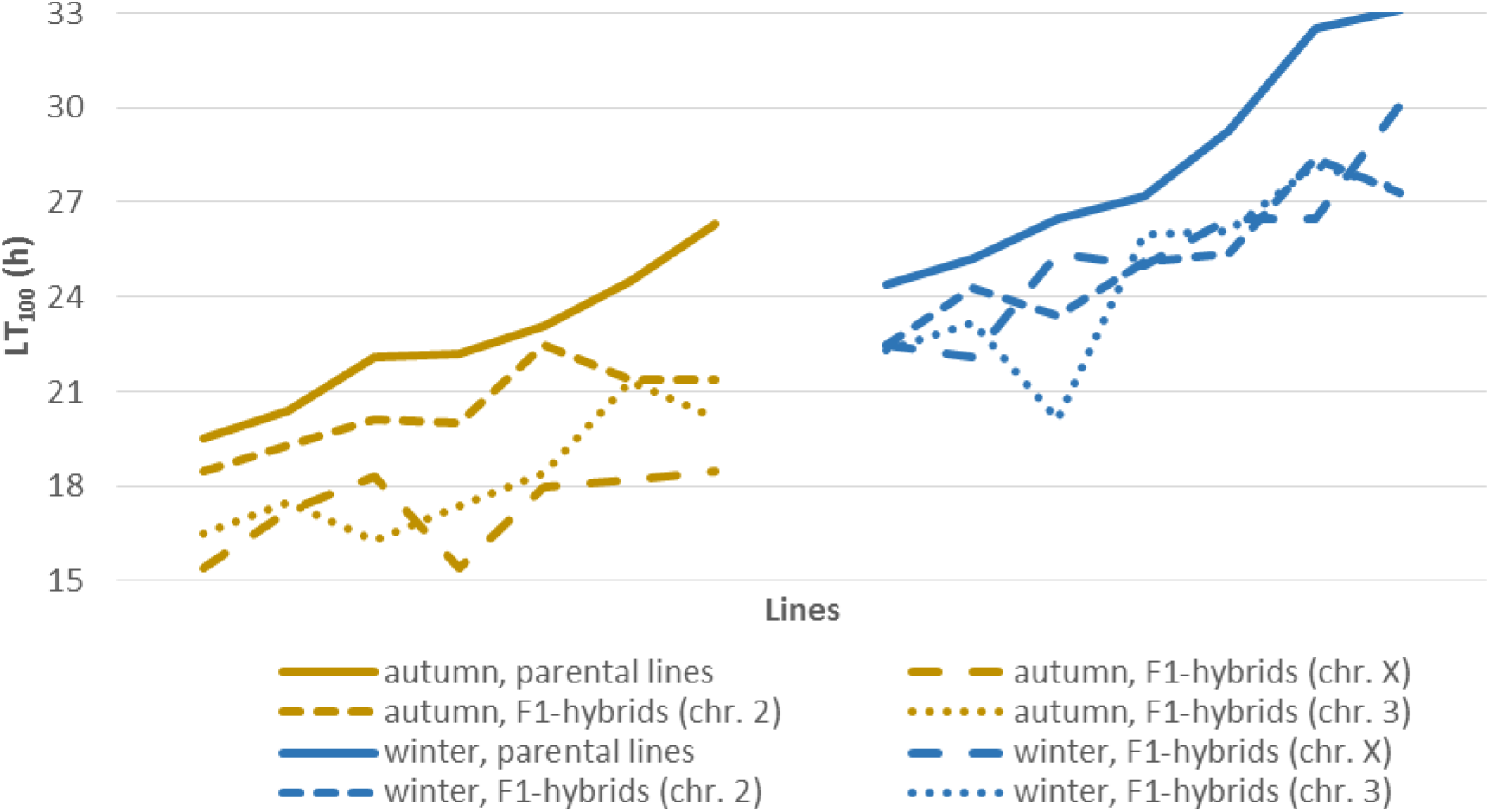
Between- and within-season variation in desiccation tolerance of the parental lines and their hybrids with three independent marker stocks.

### (b) Recombination rate

First, we compared RR in the autumn and the winter hybrids reared in normal conditions. The effects of different factors on the between-season differences in RR were estimated using a generalized linear model (see Subsection 2c). The season effect appeared significant for five intervals: one interval of chromosome X (*v–f*) and four intervals of chromosome 2 (*al–dp*, *pr–c*, *c–px* and *px–sp*). In four intervals (*v–f*, *al–dp*, *pr–c* and *c–px*), RR was higher in the winter hybrids, while one interval (*px–sp*) demonstrated the opposite pattern. The effect of season was the strongest for the pericentromeric interval *pr–c* (Table 1). The line effect was non-significant for all intervals (*p*≥0.8).

**Table 1.** The effect of season on recombination rate.

| Interval | Recombination rate (%), ± SE | | Effect of the season | | |
| --- | --- | --- | --- | --- | --- |
|  | Autumn hybrids | Winter hybrids | W | *p* | *p* (FDR) |
| Chromosome X ($N_{AC}=1820$; $N_{WC}=1797$) | | | | | |
| *y-cv* | $12.19 \pm 0.77$ | $10.96 \pm 0.73$ | 1.372 | 0.241 | 0.349 |
| *cv-v* | $19.17 \pm 0.92$ | $21.42 \pm 0.97$ | 2.837 | 0.092 | 0.200 |
| *v-f* | $19.95 \pm 0.93$ | $23.30 \pm 1.00$ | 6.135 | 0.013 | **0.045** |
| Chromosome 2 ($N_{AC}=1767$; $N_{WC}=1803$) | | | | | |
| *al-dp* | $7.35 \pm 0.62$ | $10.59 \pm 0.72$ | 11.419 | $7.3 \cdot 10^{-4}$ | **$4.7 \cdot 10^{-3}$** |
| *dp-b* | $24.90 \pm 1.02$ | $23.07 \pm 0.99$ | 1.717 | 0.190 | 0.309 |
| *b-pr* | $3.39 \pm 0.40$ | $3.94 \pm 0.46$ | 1.000 | 0.317 | 0.387 |
| *pr-c* | $14.88 \pm 0.85$ | $19.86 \pm 0.94$ | 15.422 | $8.6 \cdot 10^{-5}$ | **$1.2 \cdot 10^{-3}$** |
| *c-px* | 18.73 ± 0.94 | 21.91 ± 0.97 | 5.663 | 0.017 | **0.045** |
| *px-sp* | 5.43 ± 0.53 | 3.66 ± 0.44 | 5.933 | 0.015 | **0.045** |
| Chromosome 3 ($N_{AC}=1685$; $N_{WC}=1697$) | | | | | |
| *ru-h* | 20.36 ± 0.98 | 20.27 ± 0.98 | 0.008 | 0.930 | 0.930 |
| *h-th* | 15.25 ± 0.88 | 17.21 ± 0.91 | 2.356 | 0.125 | 0.232 |
| *th-sr* | 15.13 ± 0.87 | 15.50 ± 0.87 | 0.107 | 0.743 | 0.805 |
| *sr-e* | 6.12 ± 0.58 | 7.01 ± 0.62 | 0.960 | 0.327 | 0.387 |

$N_{AC}$ and $N_{WC}$ and $N_{WT}$ stand for the total number of flies in the autumn and winter control groups, respectively. FDR-corrected significance $p(\text{FDR}) < 0.1$ is bolded.

Second, we addressed RR plasticity to desiccation stress, by comparing RR in flies reared in normal conditions and those subjected to desiccation hardening (see Subsection 2b). We used a generalized linear model to estimate the effects of different factors on the desiccation-induced changes separately for the two seasonal cohorts and then compared these effects. We found four intervals where desiccation hardening significantly increased RR in the autumn but not winter hybrids. These include two intervals of chromosome X (*cv–v* and *v–f*) and two intervals of chromosome 2 (*pr–c* and *c–px*) (Table 2). For all these intervals, frequencies of the two reciprocal recombinants raised concordantly (not shown), which allows excluding segregation distortion as an explanation for the observed changes in RR. The line effect appeared insignificant for all intervals, in both seasonal cohorts (*p*≥0.19 and 0.46 for autumn and winter, respectively); the same holds for the line-by-treatment interaction (*p*≥0.68 and 0.49).

**Table 2.** The effects of desiccation stress on recombination rate, by seasonal cohorts.

| Interval | Season | Recombination rate (%), ± SE |  | Effect of the treatment |  |  |
| --- | --- | --- | --- | --- | --- | --- |
|  |  | Normal conditions | Desiccation-stressed | W | *p* | *P* (FDR) |
| Chromosome X (*N*<sub>AC</sub>=1820; *N*<sub>AT</sub>=1752; *N*<sub>WC</sub>=1797; *N*<sub>WT</sub>=1757) |  |  |  |  |  |  |
| *y-cv* | Autumn | 12.19 ± 0.77 | 13.13 ± 0.80 | 0.702 | 0.402 | 0.581 |
|  | Winter | 10.96 ± 0.73 | 10.02 ± 0.72 | 0.779 | 0.378 | 0.662 |
| *cv-v* | Autumn | 19.17 ± 0.92 | 23.80 ± 1.01 | 11.366 | 7.5·10<sup>-4</sup> | **0.010** |
|  | Winter | 21.42 ± 0.97 | 24.30 ± 1.03 | 4.183 | 0.041 | 0.424 |
| *v-f* | Autumn | 19.95 ± 0.93 | 23.85 ± 1.01 | 8.365 | 3.8·10<sup>-3</sup> | **0.017** |
|  | Winter | 23.30 ± 1.00 | 24.13 ± 1.02 | 0.368 | 0.544 | 0.662 |
| Chromosome 2 (*N*<sub>AC</sub>=1767; *N*<sub>AT</sub>=1716; *N*<sub>WC</sub>=1803; *N*<sub>WT</sub>=1726) |  |  |  |  |  |  |
| *al-dp* | Autumn | 7.35 ± 0.62 | 7.52 ± 0.63 | 0.042 | 0.837 | 0.837 |
|  | Winter | 10.59 ± 0.72 | 9.56 ± 0.70 | 1.088 | 0.297 | 0.662 |
| *dp-b* | Autumn | 24.90 ± 1.02 | 24.06 ± 1.03 | 0.335 | 0.563 | 0.665 |
|  | Winter | 23.07 ± 0.99 | 23.92 ± 1.03 | 0.358 | 0.550 | 0.662 |
| *b-pr* | Autumn | 3.39 ± 0.40 | 3.67 ± 0.45 | 0.212 | 0.645 | 0.699 |
|  | Winter | 3.94 ± 0.46 | 3.88 ± 0.47 | 0.029 | 0.866 | 0.912 |
| *pr-c* | Autumn | 14.88 ± 0.85 | 18.71 ± 0.94 | 9.481 | 2.1·10<sup>-3</sup> | **0.014** |
|  | Winter | 19.86 ± 0.94 | 19.99 ± 0.96 | 0.012 | 0.912 | 0.912 |
| *c-px* | Autumn | 18.73 ± 0.94 | 22.26 ± 1.00 | 6.757 | 9.3·10<sup>-3</sup> | **0.030** |
|  | Winter | 21.91 ± 0.97 | 24.33 ± 1.03 | 2.741 | 0.098 | 0.424 |
| *px-sp* | Autumn | 5.43 ± 0.53 | 4.14 ± 0.48 | 3.043 | 0.081 | 0.151 |
|  | Winter | 3.66 ± 0.44 | 4.52 ± 0.50 | 1.477 | 0.224 | 0.662 |
| Chromosome 3 ($N_{AC}=1685$; $N_{AT}=1687$; $N_{WC}=1697$; $N_{WT}=1649$) | | | | | | |
| *ru-h* | Autumn | 20.36 ± 0.98 | 21.34 ± 0.99 | 0.460 | 0.498 | 0.647 |
|  | Winter | 20.27 ± 0.98 | 19.22 ± 0.97 | 0.566 | 0.452 | 0.662 |
| *h-th* | Autumn | 15.25 ± 0.88 | 17.60 ± 0.92 | 3.259 | 0.071 | 0.151 |
|  | Winter | 17.21 ± 0.91 | 18.01 ± 0.95 | 0.340 | 0.560 | 0.662 |
| *th-sr* | Autumn | 15.13 ± 0.87 | 17.62 ± 0.92 | 3.905 | 0.048 | 0.125 |
|  | Winter | 15.50 ± 0.87 | 17.76 ± 0.94 | 2.927 | 0.087 | 0.424 |
| *sr-e* | Autumn | 6.12 ± 0.58 | 5.51 ± 0.55 | 0.707 | 0.401 | 0.581 |
|  | Winter | 7.01 ± 0.62 | 6.49 ± 0.60 | 0.425 | 0.515 | 0.662 |

$N_{AC}$ , $N_{AT}$ , $N_{WC}$ and $N_{WT}$ stand for the total number of flies in the autumn control, autumn treatment, winter control, and winter treatment groups, respectively. FDR-corrected significance $p(\text{FDR}) < 0.1$ is bolded.

### (c) Crossover interference

First, we compared the coefficients of coincidence for the autumn and the winter hybrids reared in normal conditions. The analysis was performed using an explicit maximum-likelihood model (see Subsection 2c). A consistent pattern (albeit with some exceptions mentioned below) was an increase in double-crossover rate in the winter hybrids, manifested as a relaxation of positive CI or even emergence of negative CI. Such an increase appeared significant for four pairs of intervals: two in chromosome X (*y–v*–*f* and *cv–v*–*f*) and two in chromosome 2 (*al–dp–pr* and *b–c–sp*). At that, the two former pairs represent CI in the neighborhood of the same node (marker *v*). Several pairs of intervals in chromosome 3 tended to demonstrate the opposite pattern, a decrease in double-crossover rate in the winter hybrids. For one such pair (*ru–h*–*th*), the decrease appeared significant (Table 3) The line effect (within-season heterogeneity) was significant for several pairs of intervals in chromosome 2, only in the autumn but not winter hybrids; all such pairs represent CI in the neighborhood of two nodes (markers *dp* and *c*). For all other pairs, both seasonal cohorts were homogenous (not shown).

**Table 3.**
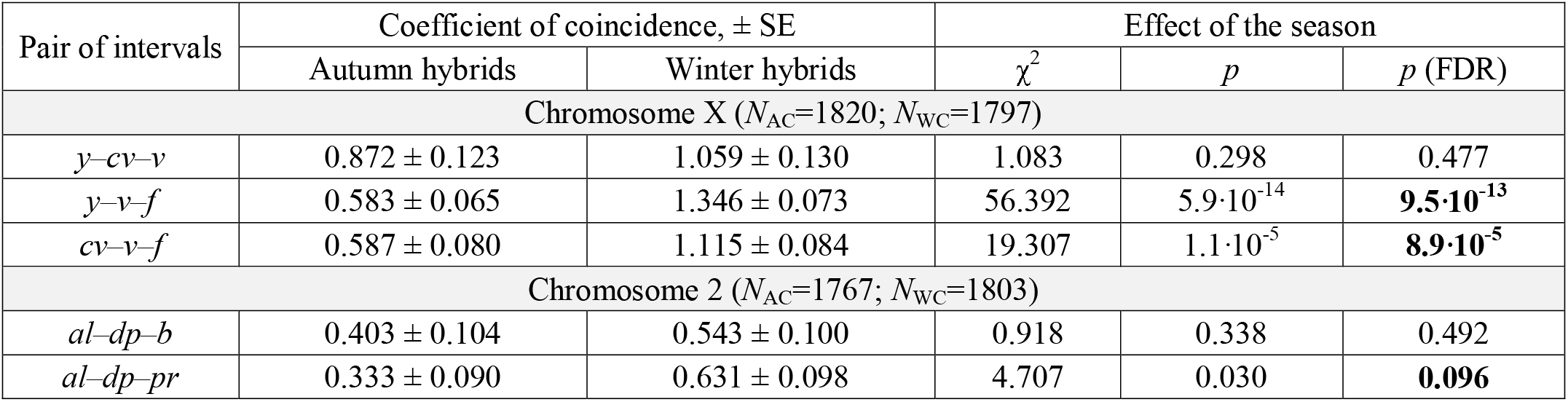
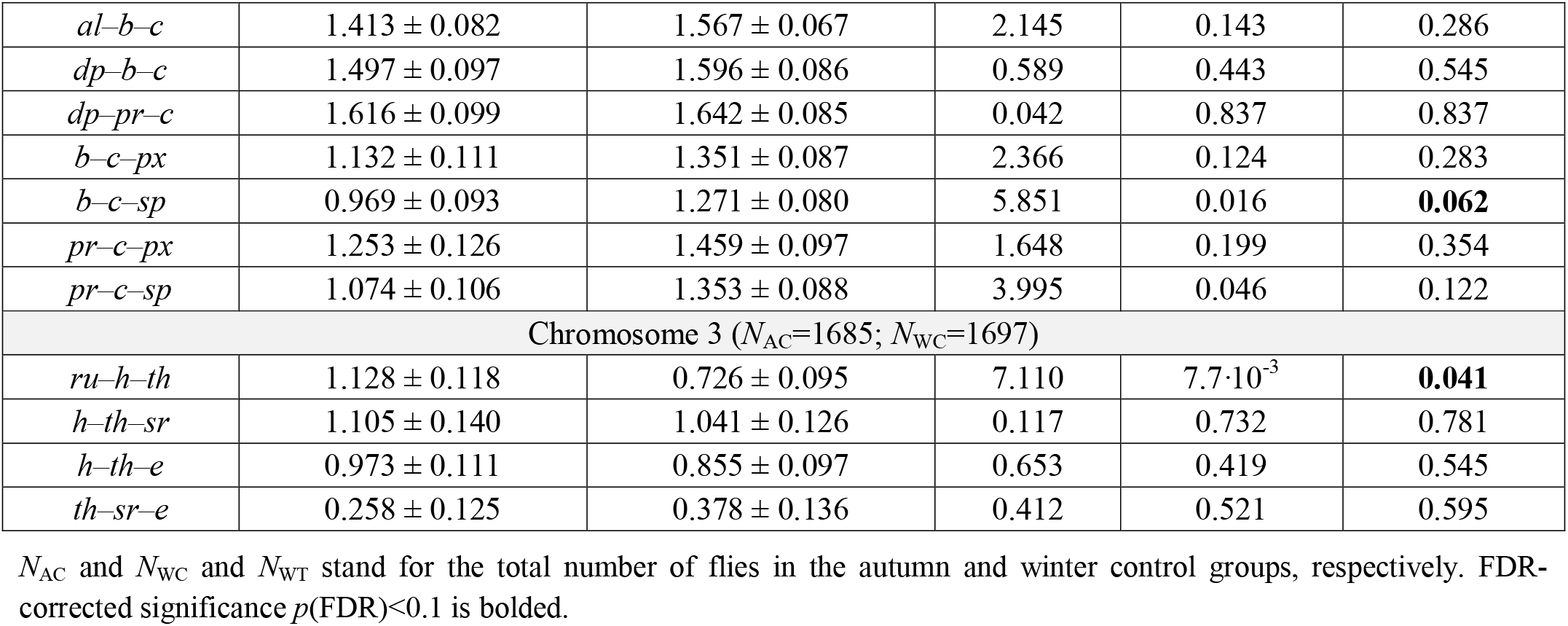
The effect of season on crossover interference.

| Pair of intervals | Coefficient of coincidence, ± SE |  | Effect of the season |  |  |
| --- | --- | --- | --- | --- | --- |
| | Autumn hybrids | Winter hybrids | $\chi^2$ | $p$ | $p$ (FDR) |
| Chromosome X ($N_{AC}=1820$; $N_{WC}=1797$) | | | | | |
| $y-cv-v$ | 0.872 ± 0.123 | 1.059 ± 0.130 | 1.083 | 0.298 | 0.477 |
| $y-v-f$ | 0.583 ± 0.065 | 1.346 ± 0.073 | 56.392 | $5.9 \cdot 10^{-14}$ | **$9.5 \cdot 10^{-13}$** |
| $cv-v-f$ | 0.587 ± 0.080 | 1.115 ± 0.084 | 19.307 | $1.1 \cdot 10^{-5}$ | **$8.9 \cdot 10^{-5}$** |
| Chromosome 2 ($N_{AC}=1767$; $N_{WC}=1803$) | | | | | |
| $al-dp-b$ | 0.403 ± 0.104 | 0.543 ± 0.100 | 0.918 | 0.338 | 0.492 |
| $al-dp-pr$ | 0.333 ± 0.090 | 0.631 ± 0.098 | 4.707 | 0.030 | **0.096** |
| *al-b-c* | 1.413 ± 0.082 | 1.567 ± 0.067 | 2.145 | 0.143 | 0.286 |
| *dp-b-c* | 1.497 ± 0.097 | 1.596 ± 0.086 | 0.589 | 0.443 | 0.545 |
| *dp-pr-c* | 1.616 ± 0.099 | 1.642 ± 0.085 | 0.042 | 0.837 | 0.837 |
| *b-c-px* | 1.132 ± 0.111 | 1.351 ± 0.087 | 2.366 | 0.124 | 0.283 |
| *b-c-sp* | 0.969 ± 0.093 | 1.271 ± 0.080 | 5.851 | 0.016 | **0.062** |
| *pr-c-px* | 1.253 ± 0.126 | 1.459 ± 0.097 | 1.648 | 0.199 | 0.354 |
| *pr-c-sp* | 1.074 ± 0.106 | 1.353 ± 0.088 | 3.995 | 0.046 | 0.122 |
| Chromosome 3 ($N_{AC}=1685$; $N_{WC}=1697$) | | | | | |
| *ru-h-th* | 1.128 ± 0.118 | 0.726 ± 0.095 | 7.110 | $7.7 \cdot 10^{-3}$ | **0.041** |
| *h-th-sr* | 1.105 ± 0.140 | 1.041 ± 0.126 | 0.117 | 0.732 | 0.781 |
| *h-th-e* | 0.973 ± 0.111 | 0.855 ± 0.097 | 0.653 | 0.419 | 0.545 |
| *th-sr-e* | 0.258 ± 0.125 | 0.378 ± 0.136 | 0.412 | 0.521 | 0.595 |

$N_{AC}$ and $N_{WC}$ and $N_{WT}$ stand for the total number of flies in the autumn and winter control groups, respectively. FDR-corrected significance $p(\text{FDR}) < 0.1$ is bolded.

Second, we compared the coefficients of coincidence for flies reared in normal conditions and flies subjected to desiccation hardening. The effect of treatment was estimated using an explicit maximum-likelihood model (see Subsection 2c); such analysis was conducted separately for each of the two seasonal cohorts, and then the revealed effects were compared. Desiccation hardening tended to increase the double-crossover rate in the autumn hybrids and decrease it in the winter hybrids, but the effect appeared significant only for some pairs of intervals. For three interval pairs in chromosome X (*y–cv–v*, *y–v–f* and *cv–v–f*) and two in chromosome 2 (*al–dp–pr* and *al–b–c*), the autumn hybrids demonstrated a significant increase in double-crossover rate upon desiccation hardening, while their winter counterparts did not respond significantly to the treatment. For another pair of intervals in chromosome 3 (*ru–h–th*), the two seasonal cohorts behaved oppositely: a significant increase in double-crossover rate was observed only in the winter but not autumn hybrids (Table 4).

**Table 4.** The effects of desiccation stress on crossover interference, by seasonal cohorts.

| Pair of intervals | Season | Coefficient of coincidence, ± SE | | Effect of the treatment | | |
| --- | --- | --- | --- | --- | --- | --- |
| | | Normal Conditions | Desiccation-stressed | Z | $p$ | $p$ (FDR) |
| Chromosome X ($N_{AC}$=1820; $N_{AT}$=1752; $N_{WC}$=1797; $N_{WT}$=1757) | | | | | | |
| $y$-$cv$-$v$ | Autumn | $0.872 \pm 0.123$ | $1.207 \pm 0.116$ | 1.928 | 0.054 | 0.172 |
| | Winter | $1.059 \pm 0.130$ | $1.214 \pm 0.133$ | 0.793 | 0.427 | 0.714 |
| $y$-$v$-$f$ | Autumn | $0.583 \pm 0.065$ | $1.173 \pm 0.068$ | 5.937 | $2.9 \cdot 10^{-9}$ | **$4.6 \cdot 10^{-8}$** |
| | Winter | $1.346 \pm 0.073$ | $1.095 \pm 0.068$ | -2.220 | 0.026 | 0.211 |
| $cv$-$v$-$f$ | Autumn | $0.587 \pm 0.080$ | $1.047 \pm 0.077$ | 3.894 | $9.9 \cdot 10^{-5}$ | **$7.9 \cdot 10^{-4}$** |
| | Winter | $1.115 \pm 0.084$ | $1.008 \pm 0.075$ | -0.801 | 0.423 | 0.714 |
| Chromosome 2 ($N_{AC}$=1767; $N_{AT}$=1716; $N_{WC}$=1803; $N_{WT}$=1726) | | | | | | |
| $al$-$dp$-$b$ | Autumn | $0.403 \pm 0.104$ | $0.605 \pm 0.110$ | 1.495 | 0.135 | 0.308 |
| | Winter | $0.543 \pm 0.100$ | $0.738 \pm 0.135$ | 0.737 | 0.461 | 0.714 |
| *al-dp-pr* | Autumn | $0.333 \pm 0.090$ | $0.672 \pm 0.107$ | 3.430 | $6.0 \cdot 10^{-4}$ | **$3.2 \cdot 10^{-3}$** |
| | Winter | $0.631 \pm 0.098$ | $0.909 \pm 0.134$ | 0.679 | 0.497 | 0.714 |
| *al-b-c* | Autumn | $1.413 \pm 0.082$ | $1.654 \pm 0.069$ | 3.197 | $1.4 \cdot 10^{-3}$ | **$5.6 \cdot 10^{-3}$** |
| | Winter | $1.567 \pm 0.067$ | $1.761 \pm 0.075$ | 0.758 | 0.449 | 0.714 |
| *dp-b-c* | Autumn | $1.497 \pm 0.097$ | $1.640 \pm 0.085$ | 1.652 | 0.099 | 0.263 |
| | Winter | $1.596 \pm 0.086$ | $1.698 \pm 0.088$ | 0.415 | 0.678 | 0.750 |
| *dp-pr-c* | Autumn | $1.616 \pm 0.099$ | $1.679 \pm 0.085$ | 1.187 | 0.235 | 0.376 |
| | Winter | $1.642 \pm 0.085$ | $1.755 \pm 0.089$ | 0.381 | 0.703 | 0.750 |
| *b-c-px* | Autumn | $1.132 \pm 0.111$ | $1.423 \pm 0.083$ | 0.152 | 0.879 | 0.879 |
| | Winter | $1.351 \pm 0.087$ | $1.169 \pm 0.089$ | 0.557 | 0.578 | 0.714 |
| *b-c-sp* | Autumn | $0.969 \pm 0.093$ | $1.358 \pm 0.077$ | 0.810 | 0.418 | 0.557 |
| | Winter | $1.271 \pm 0.080$ | $1.089 \pm 0.080$ | 0.816 | 0.415 | 0.714 |
| *pr-c-px* | Autumn | $1.253 \pm 0.126$ | $1.492 \pm 0.092$ | -0.240 | 0.810 | 0.864 |
| | Winter | $1.459 \pm 0.097$ | $1.237 \pm 0.099$ | 0.145 | 0.885 | 0.885 |
| *pr-c-sp* | Autumn | $1.074 \pm 0.106$ | $1.431 \pm 0.085$ | 0.397 | 0.691 | 0.790 |
| | Winter | $1.353 \pm 0.088$ | $1.157 \pm 0.090$ | 0.615 | 0.538 | 0.714 |
| Chromosome 3 ($N_A=3372$; $N_W=3346$) | | | | | | |
| *ru-h-th* | Autumn | $1.128 \pm 0.118$ | $1.183 \pm 0.106$ | 0.400 | 0.689 | 0.790 |
| | Winter | $0.726 \pm 0.095$ | $1.190 \pm 0.113$ | 3.230 | $1.2 \cdot 10^{-3}$ | **$2.0 \cdot 10^{-2}$** |
| *h-th-sr* | Autumn | $1.105 \pm 0.140$ | $0.877 \pm 0.109$ | -1.277 | 0.202 | 0.376 |
| | Winter | $1.041 \pm 0.126$ | $1.122 \pm 0.117$ | 0.553 | 0.580 | 0.714 |
| *h-th-e* | Autumn | $0.973 \pm 0.111$ | $0.799 \pm 0.092$ | -1.195 | 0.232 | 0.376 |
| | Winter | $0.855 \pm 0.097$ | $0.950 \pm 0.094$ | 0.741 | 0.459 | 0.714 |
| *th-sr-e* | Autumn | $0.258 \pm 0.125$ | $0.426 \pm 0.153$ | 1.074 | 0.283 | 0.412 |
| | Winter | $0.378 \pm 0.136$ | $0.374 \pm 0.135$ | 0.681 | 0.496 | 0.714 |

$N_{AC}$ , $N_{AT}$ , $N_{WC}$ and $N_{WT}$ stand for the total number of flies in the autumn control, autumn treatment, winter control, and winter treatment groups, respectively. FDR-corrected significance $p(\text{FDR}) < 0.1$ is bolded.

## 4. Discussion

### (a) Seasonal changes in desiccation tolerance

The winter parental lines demonstrated significantly higher desiccation tolerance than the autumn lines. Remarkably, this difference held also for all three types of hybrids, obtained by crossing the parental wild type lines with the three marker stocks. Moreover, all four estimates of desiccation tolerance (measured in the parental lines and the three hybrids) significantly correlated (Fig. 1). These findings, together with the fact that the parental lines were kept over five to six generations in normal conditions before the desiccation-tolerance assay, strongly argue for genetic rather than plastic nature of the observed changes in stress tolerance. At that, the underlying desiccation-conferring alleles should be non-recessive, implying that natural selection should be strong enough to cause genetic adaptation during just ~8-10 generations (about five times shorter than our previous artificial-selection experiment [41]). Such a quick response to selection pressure confirms the conclusion of Hoffmann and Harshman [42] that *D. melanogaster* has very high heritability of desiccation tolerance, perhaps unusually high for the whole genus.

### (b) Seasonal changes in recombination rate

In five intervals (*v–f*, *al–dp*, *pr–c*, *c–px* and *px–sp*), RR significantly differed between the seasons. These genome regions remarkably overlap with those that had demonstrated a considerable correlated recombination response after 48 generations of artificial directional selection for desiccation tolerance in our previous study [20]. In both cases, we consider the observed changes in RR to be genetic (i.e., caused by microevolutionary changes in recombination-modifying alleles) rather than plastic. The precondition for such changes, the presence of standing variation in recombination-modifying genes, holds for natural *D. melanogaster* populations, as suggested by several classical experiments that had succeeded to select for both decreased and increased RR [43–45]. The question, however, is which forces drive recombination evolution under *indirect* selection for RR, like in the current study or in those experiments where recombination coevolved along with other, recombination-*unrelated* traits [20–22,46–51].

Theoretical models suggest that indirect selection on increased RR may arise from directional selection on a recombination-unrelated fitness trait. Local episodes of directional selection are relatively common in nature and are believed to be a powerful force in recombination evolution [52–54]. Within the ‘indirect-selection’ explanation, two different, although non-excluding, mechanisms are discussed in the literature. According to the first mechanism, if epistasis between the trait-affecting loci is negative, then negative linkage disequilibria (LD) will tend to emerge in the population favouring increased RR. This mechanism works in large populations, especially of species with a small number of chromosomes [55]. However, later analysis has shown that such a long-term advantage of increased recombination may sometimes be outbalanced by its short-term disadvantage. It turned out that directional selection on a fitness-related trait may favour increased RR only if negative epistasis is not too strong or, otherwise, if the recombination-modifying locus is tightly linked to the trait-affecting loci [56]. According to the second mechanism, LD between the trait-affecting loci are permanently generated by drift, and their sign is thus determined by chance. Yet, selection eliminates the positive LD more easily, and the prevailing negative LD again favour increased RR. This mechanism should work in small to moderate-size populations, even when the fitness-related trait is controlled by purely multiplicative genes [57,58].

As discussed in the previous subsection, the observed seasonal changes in desiccation tolerance seem to be mostly inheritable and reflect short-term but powerful episodes of natural selection. This makes the indirect-selection explanation reasonable, although specific mechanisms within this explanation (negative epistasis or drift) cannot be inferred. Besides, it must be mentioned that desiccation was not a single environmental stressor. During the winter, the population had to cope also with low temperature, low food availability, and probably higher pathogen prevalence. Previous studies have shown that resistance to temperature, starvation, and infections also oscillates across the year in natural *D. melanogaster* populations. Often, these seasonal changes reflect genetic adaptation rather than plasticity [15–17] (but see [59,60] for the opposite findings). Thus, the presumed episode of natural selection was surely more complex than those of artificial selection previously created in the lab. Such multidimensional selection might further favour increased RR, given the recognized potential of recombination to mitigate selection slowing caused by the Hill-Robertson effect [61,62].

Several alternatives to the indirect-selection explanation exist. First, recombination-increasing alleles might initially be linked, by chance, to the favourable selected alleles or haplotypes. An implicit objection to this comes from the fact that increase in RR in earlier studies was observed even when selection acted in two opposite directions (early/late flowering time [22], positive/negative geotaxis [50], sternopleural bristle number [51] or hypoxia/hyperoxia tolerance [20]). The observed between-season differences in RR may also result from random changes in frequencies of recombination-modifying alleles. This scenario, however, seems less realistic given the remarkable concordance of the results within each season (the line effect was non-significant). A similar concordance was also observed in our previous artificial-selection experiment [20]. Finally, the differences may be associated with epigenetic changes in recombination that have succeeded to pass through generations. Indeed, several studies have reported changes in RR in untreated individuals whose parents and even grandparents were subjected to a stressor [38,63–65]. Yet, it seems unrealistic that such ‘epigenetic memory’ had lasted for five-six generations during which isofemale lines were kept in the lab under normal conditions before the recombination-assessment experiment.

### (c) Desiccation-induced changes in recombination rate and their modulation by season and desiccation tolerance

In five intervals (*cv–v*, *v–f*, *pr–c*, *c–px*, and *th–sr*), RR significantly increased in flies exposed to desiccation hardening, compared to flies with the same genotype but reared in normal conditions. This result contributes to extremely scarce evidence for the recombinogenic effect of desiccation [20,66]. Such an effect can be explained mechanistically, via biochemical effects of desiccation on recombination enzymes. Water deficit raises the intracellular concentration of reactive oxygen species, well-recognized DNA-damaging agents [67,68]. The elevated level of DNA damages under dehydration has been reported for different species, including Diptera [69–71]. The damage-induced DNA repair may also intensify recombination due to the upregulation of some common enzymatic pathways [72,73]. Additionally, the increased RR in desiccation-stressed flies may be associated with changes in the ‘recombination-reacting’ system via gene expression, which naturally requires chromatin unfolding, increases DNA accessibility to recombination machinery [74,75]. As suggested by the gene-ontology analysis, the chromosome regions where RR has plastically increased upon desiccation hardening are indeed significantly enriched by genes related to desiccation tolerance (Table S1). These include genes whose products are involved in oxidation/reduction processes (intervals *pr–c* and *th–sr*) and stress-signalling (interval *c–px*). This result is in line with the well-recognized role of oxidation/reduction in the acute adaptive response to desiccation stress, including in *D. melanogaster* [76,77]. Remarkably, genome regions closed to intervals *pr–c* and *th–sr* demonstrated pronounced RR plasticity also to heat [26]. Besides, interval *c–px* also harbours many genes affecting the catabolism of chitin, which is an important biochemical pathway for water-loss control in insects [78].

Importantly, the observed RR response to the desiccation hardening significantly differed for the two seasonal cohorts, being much higher in the autumn than the winter flies (even despite a longer exposure time for the latter). This suggests that stressor-induced changes in RR may be modulated by the severity of stress experienced by an organism rather than the value of the stressor itself. Such a negative association between RR plasticity and stress-tolerance was observed in several empirical studies [18,79–81], including our recent experiment with desiccation-treated fruit flies [28]. Moreover, theoretical models have demonstrated that this association may be advantageous in a changing environment and evolve as an adaptive strategy [82,83]. The season-specific RR response to desiccation hardening revealed in the current study has urged us to explore the plasticity-fitness association in more detail. We found that in most cases, an increase in RR is negatively correlated with the genotype’s desiccation tolerance. The association was stronger for desiccation tolerance of the hybrid (compared to that of the parental line), and for the relative increase in RR (compared to absolute one) (Table S2).

### (d) Seasonal and desiccation-induced changes in crossover interference

The seasonal changes in CI appeared significant for five pairs of intervals (*y–v–f*, *cv–v–f*, *al–dp–pr*, *b–c–sp* and *ru–h–th*). Moreover, in two former pairs in chromosomes X the double-crossover rate increased in the winter hybrids to such extent that the initial positive CI had turned into significant negative. We observed the same tendency, an increase in double-crossover rate, also for other pairs, although there it was insignificant. Remarkably, the pair of intervals *cv–v–f* in chromosome X was highly reactive also in our previous study: there, negative CI had emerged upon artificial selection, not only for desiccation tolerance but also for resistance to hypoxia and hyperoxia [20]. At the same time, several other pairs that were reactive in that evolutionary experiment (e.g., *y–cv–v*) did not show significant seasonal changes now, in the studied natural population. An interesting finding is the pair of intervals *ru–h–th* in chromosome 3, where the double-crossover rate significantly decreased in the winter hybrids. A plausible explanation for the less pronounced changes in CI in nature (compared to those in the lab) is a much shorter period of selection (~8–10 vs ~50–200 generations). Moreover, the initially high double-crossover rate may be an important factor in the autumn cohort. Such negative CI, observed in many regions (especially in chromosome 2), might have resulted from an unknown recent episode of strong selection, and probably hampered further evolution of this recombination feature. The presence of negative CI in natural populations or upon disturbances caused by environmental or genomic stresses is debated [84–87], and our finding is an important contribution to the limited empirical evidence for this phenomenon.

The desiccation hardening caused significant changes in CI in five pairs of intervals (*y–v–f*, *cv–v–f*, *al–dp–pr*, *al–b–c* and *ru–h–th*), in all three chromosomes. Usually, even when the observed changes were insignificant, the treatment tended to increase double-crossover rate. This finding is consistent with the results of other studies where an increase in double-crossover rate was observed under heat [24–26,88], hypoxia and hyperoxia [20], or even intrinsic genome stress caused by a meiosis-deregulating mutations [89–92].

Like in the case of RR, the autumn hybrids in our study tended to respond to the desiccation hardening more pronouncedly than their winter counterparts (except for pair *ru–h–th* with a significant opposite pattern). It is natural to assume that the treatment caused a more severe physiological stress to the autumn hybrids, in support of the above-discussed concept of the negative fitness-dependent control of recombination. Again, we tested for the possible association between CI plasticity to desiccation and the genotype’s desiccation tolerance. The correlation appeared negative for many pairs of intervals (all pairs in chromosome X and some of the pairs of chromosomes 2 and 3). For all pairs representing CI in the neighbourhood of marker *c* (chromosome 2) the correlation appeared positive. Usually, the association was stronger with desiccation tolerance of the hybrids rather than parental lines (Table S3).

### (e) Functional relevance of the recombination-reactive intervals

In their pioneering study, Flexon and Rodell [47] revealed a certain correlation between the recombination response to indirect selection in a given genome region and its involvement in stress tolerance. To test for such association, we performed gene-enrichment analysis. It turned out that genome regions where RR significantly differed between the seasons are enriched with genes related to desiccation tolerance and general stress response (Table S1). Thus, interval *v-f* (X chromosome) includes genes involved in signal transduction pathways and oxidoreductase activity. The role of oxidation/reduction in the acute adaptive response to desiccation stress is well-recognized [68,93,94]. In chromosome 2, interval (*al*–*dp*) is enriched by membrane formation and transmembrane transport genes; interval (*pr–c*) - by genes involved in oxidation-reduction process, cell surface receptor signaling pathway, and alpha-amilase domain involved in carbohydrate metabolic process; (*c–px*) – by stress signaling; (*px-sp*) - by genes participating in cellular water homeostasis, “major intrinsic protein [a family that form transmembrane channels which facilitate diffusion of water], and aquaporins known as water channels and considered to be the cells’ “plumbing” system [95]. The pair of the adjacent intervals *ru–h–th* in chromocome 3 includes several categories involved in seasonal adaptation and desiccation stress resistance: structural constituent of cuticle and chitin-based cuticle development, chitin metabolic process, and heat shock protein domains. This pair of intervals demonstrated a decrease in double-crossover rate in the winter hybrids (Table 3) and increase upon desiccation hardening (Table 4). The revealed correspondence between the functional effects of the tested intervals and seasonal changes in their RR and CI, as well as in plastic response to stress, calls for further studies for a more detailed characterizing the phenomenon of seasonal variation of recombination and for uncovering the underlying mechanisms.

## 5. Conclusions

Organisms in seasonal environments must integrate information from multiple environmental cues to synchronize transitions between life-history stages, imposing direct and indirect selection on multiple fitness-related traits and functions. We demonstrate that these traits also include recombination rate, crossover interference, and their plasticity to desiccation stress in a *D. melanogaster* population subject to humid autumns and dry winters in India. Thus, this study provides new evidence for seasonal variation in recombination and its plasticity in natural populations, indirectly driven by selection on seasonality-associated stress resistance. Specifically, plasticity of both recombination rate and crossover interference to desiccation after summer tends to be more pronounced than in winter. Notably, these changes accumulate within a short period (8–10 generations) of adaptation to a natural seasonal stressor.

## Acknowledgments

SR is deeply thankful to Nataliya Rybnikova for her assistance in surveying the literature and processing the data.

