## Supplementary material for "Seasonal changes in recombination rate, crossover interference, and their response to desiccation stress in a natural population of *Drosophila melanogaster* from India": Table S1

**Table S1.** Gene Ontology enrichment tests for genes in the tested intervals of chromosomes X, 2 and 3 (contrasted with all *D. melanogaster* genes using DAVID for functional enrichment analysis

| Category | Interval | Term | Gene Count | *p*-value | Fold enrichment | *p-*corrected (Benjamini-Hochberg) |
| --- | --- | --- | --- | --- | --- | --- |
|  |  | **X Chromosome** |  |  |  |  |
| UP_KEYWORDS | ***y-cv*** | Polymorphism | 18 | 4.63E-07 | 4.5 | 8.57E-05 |
| UP_KEYWORDS |  | Coiled coil | 79 | 2.98E-04 | 1.5 | 2.70E-02 |
| UP_KEYWORDS |  | Neurogenesis | 10 | 2.0E-03 | 3.6 | 1.19E-01 |
| UP_KEYWORDS |  | RNA-binding | 15 | 3.10E-03 | 2.5 | 1.36E-01 |
| UP_KEYWORDS | ***cv-v*** | Coiled coil | 87 | 9.41E-05 | 1.6 | 1.80E-02 |
| INTERPRO |  | Biotinidase, eukaryotic | 4 | 1.74E-04 | 28.1 | 1.16E-01 |
| GOTERM_MF_DIRECT | ***v-f*** | Protein kinase regulator activity | 14 | 2.92E-12 | 13.1 | 1.20E-09 |
| INTERPRO |  | **Glucose-methanol-choline oxidoreductase** | 10 | 6.85E-10 | 16.2 | 1.53E-07 |
| GOTERM_MF_DIRECT |  | **Choline dehydrogenase activity** | 7 | 6.51E-08 | 21.5 | 8.96E-06 |
| GOTERM_MF_DIRECT |  | **Oxidoreductase activity** | 11 | 1.64E-08 | 10.8 | 3.39E-06 |
| GOTERM_BP_DIRECT |  | **Signal transduction** | 26 | 8.27E-08 | 3.6 | 2.88E-05 |
| KEGG_PATHWAY |  | **Wnt signaling pathway** | 18 | 2.76E-06 | 3.8 | 2.23E-04 |
| GOTERM_BP_DIRECT |  | Protein phosphorylation | 28 | 3.71E-05 | 2.4 | 9.70E-03 |
|  |  | **2^nd^ Chromosome** |  |  |  |  |
| INTERPRO | ***al-dp*** | Proteinase inhibitor I2 | 23 | 3.21E-20 | 13.8 | 2.06E-17 |
| GOTERM_MF_DIRECT |  | Endopeptidase inhibitor activity | 23 | 1.72E-13 | 7. 4 | 5.64E-11 |
| INTERPRO |  | C-type lectin-fold | 10 | 4.61E-05 | 5.7 | 7.37E-03 |
| GOTERM_BP_DIRECT |  | Periodic partitioning by pair rule gene | 6 | 1.98E-05 | 15.8 | 1.68E-02 |
| GOTERM_CC_DIRECT |  | **Membrane** | 29 | 2.69E-04 | 2.1 | 5.76E-02 |
| GOTERM_BP_DIRECT |  | **Glucose import** | 6 | 3.15E-04 | 9.5 | 8.59E-02 |
| GOTERM_MF_DIRECT |  | **Carbohydrate binding** | 12 | 3.78E-04 | 3.7 | 4.05E-02 |
| GOTERM_MF_DIRECT |  | **Hydrolase activity** | 19 | 4.89E-04 | 2.5 | 3.93E-02 |
| GOTERM_MF_DIRECT |  | **Glucose transmembrane transporter activity** | 6 | 7.76E-04 | 7.8 | 4.97E-02 |
| UP_SEQ_FEATURE |  | **Transmembrane region** | 33 | 3.33E-04 | 1.8 | 1.02E-01 |
| COG_ONTOLOGY | ***dp-b*** | Lipid metabolism | 13 | 7.41E-04 | 3.0 | 1.84E-02 |
| UP_KEYWORDS |  | Extracellular matrix | 9 | 2.19E-04 | 5.0 | 5.75E-02 |
| GOTERM_MF_DIRECT |  | **Carbohydrate binding** | 17 | 1.45E-03 | 2.4 | 2.11E-01 |
| INTERPRO |  | **Glycoside hydrolase family** | 6 | 1.64E-04 | 9.2 | 1.00E-01 |
| KEGG_PATHWAY | ***b-pr*** | Retinol metabolism | 7 | 7.75E-05 | 9.1 | 5.40E-03 |
| PIR_SUPERFAMILY |  | **Short chain dehydrogenase** | 4 | 7.81E-05 | 22.2 | 9.30E-03 |
| UP_KEYWORDS |  | Metalloprotease | 10 | 4.46E-05 | 5.8 | 9.68E-03 |
| INTERPRO |  | Peptidase M12A, astacin | 6 | 4.46E-05 | 13.6 | 2.73E-02 |
| SMART |  | ZnMc | 6 | 5.02E-05 | 12.3 | 1.06E-02 |
| SMART | ***pr-c*** | **Alpha-amylase domain (carbohydrate metabolic process)** | 12 | 2.36E-08 | 6.9 | 8.18E-06 |
| INTERPRO |  | **Glycoside hydrolase** | 12 | 2.33E-02 | 6.2 | 2.31E-04 |
| GOTERM_BP_DIRECT |  | **Cell surface receptor signaling pathway** | 24 | 1.33E-07 | 3.2 | 6.40E-04 |
| GOTERM_MF_DIRECT |  | **Monooxygenase activity** | 28 | 3.53E-07 | 2.6 | 2.08E-03 |
| GOTERM_MF_DIRECT |  | **Oxidoreductase activity** | 27 | 1.42E-05 | 2.4 | 4.41E-03 |
| KEGG_PATHWAY |  | Starch and sucrose metabolism | 14 | 5.75E-05 | 3.5 | 6.70E-03 |
| UP_SEQ_FEATURE |  | Metal ion-binding | 27 | 1.42E-05 | 2.3 | 2.92E-02 |
| GOTERM_CC_DIRECT |  | **Organelle membrane** | 24 | 3.29E-05 | 2.5 | 2.50E-02 |
| GOTERM_MF_DIRECT |  | **Substrate-specific transmembrane transporter activity** | 11 | 5.27E-05 | 4.3 | 1.20E-02 |
| GOTERM_CC_DIRECT |  | **Extracellular space** | 104 | 6.55E-05 | 1.4 | 2.44E-02 |
| GOTERM_BP_DIRECT |  | **Sensory perception of chemical stimulus** | 25 | 7.41E-05 | 2.4 | 6.50E-02 |
| GOTERM_MF_DIRECT |  | Odorant binding | 27 | 7.19E-04 | 2.0 | 7.31E-02 |
| GOTERM_BP_DIRECT |  | **Oxidation-reduction process** | 71 | 1.03E-04 | 1.5 | 1.69E-01 |
| UP_KEYWORDS |  | **Oxidoreductase** | 98 | 5.81E-04 | 1.3 | 2.07E-01 |
| UP_KEYWORDS | ***c-px*** | **Signal transduction** | 98 | 6.32E-04 | 1.3 | 1.40E-01 |
| SMART |  | LRR_TYP | 8 | 2.47E-03 | 4.2 | 1.95E-01 |
| SMART | ***px-sp*** | LITAF lipopolysaccharide induced TNF factor | 13 | 4.14E-15 | 24.5 | 4.93E-13 |
| INTERPRO |  | LPS-induced tumor necrosis alpha factor | 13 | 1.76E-02 | 24.9 | 2.42E-12 |
| GOTERM_MF_DIRECT |  | **Water channel activity** | 4 | 1.05E-03 | 18.1 | 1.45E-01 |
| GOTERM_MF_DIRECT |  | Glycerol channel activity | 4 | 4.54E-15 | 20.6 | 1.82E-01 |
| INTERPRO |  | **Major intrinsic protein** | 4 | 7.72E-04 | 19.7 | 1.86E-01 |
| INTERPRO |  | **Aquaporin-like** | 4 | 1.21E-03 | 17.3 | 1.93E-01 |
|  |  | **3^rd^ Chromosome** |  |  |  |  |
| INTERPRO | ***ru-h*** | **Insect cuticle protein** | 36 | 1.70E-16 | 5.3 | 2.03E-13 |
| GOTERM_MF_DIRECT |  | **Structural constituent of chitin-based larval cuticle** | 34 | 4.02E-15 | 5.1 | 1.75E-12 |
| GOTERM_MF_DIRECT |  | **Structural constituent of cuticle** | 34 | 2.41E-14 | 4.8 | 5.29E-12 |
| GOTERM_BP_DIREC |  | **Chitin-based cuticle development** | 37 | 7.74E-13 | 4.0 | 9.08E-10 |
| GOTERM_CC_DIRECT |  | Extracellular matrix | 35 | 6.34E-12 | 3.9 | 1.88E-09 |
| INTERPRO |  | Gustatory receptor | 6 | 4.64E-05 | 11.9 | 2.10E-02 |
| SMART |  | Zn_pept | 8 | 1.59E-04 | 6.3 | 1.59E-02 |
| GOTERM_MF_DIREC |  | Metallo-carboxypeptidase  activity | 8 | 3.88E-04 | 5.5 | 3.35E-02 |
| GOTERM_BP_DIRECT |  | Detection of chemical stimulus involved in sensory perception of taste | 8 | 8.29E-05 | 6.9 | 4.74E-02 |
| GOTERM_BP_DIRECT |  | Sensory perception of sweet taste | 6 | 5.52E-04 | 8.0 | 1.94E-01 |
| GOTERM_BP_DIRECT | ***h-th*** | **Chitin metabolic process** | 30 | 3.26E-15 | 6.0 | 3.50E-12 |
| INTERPRO |  | **Chitin binding domain** | 30 | 2.01E-13 | 5.2 | 1.75E-10 |
| GOTERM_CC_DIRECT |  | Extracellular region | 63 | 4.11E-08 | 2.1 | 1.33E-05 |
| INTERPRO |  | **Alpha crystallin/Hsp20 domain** | 7 | 3.17E-05 | 9.9 | 9.14 E-03 |
| INTERPRO |  | **Alpha crystallin/Heat shock protein** | 6 | 4.98E-05 | 12.2 | 8.62E-03 |
| INTERPRO |  | Insulin family | 5 | 4.19E-05 | 18.3 | 9.07E-03 |
| GOTERM_BP_DIRECT |  | Defense response to bacterium | 14 | 2.33E-05 | 4.2 | 1.25E-02 |
| INTERPRO |  | **HSP20-like chaperone** | 7 | 9.03E-04 | 5.8 | 1.06 E-01 |
| GOTERM_CC_DIRECT | ***th-sr*** | **Integral component of plasma membrane** | 129 | 1.53E-07 | 1.5 | 8.11E-05 |
| GOTERM_MF_DIRECT |  | Zinc ion binding | 146 | 1.77E-04 | 1.3 | 8.16E-02 |
| GOTERM_BP_DIRECT |  | **Regulation of transcription, DNA-templated** | 94 | 1.09E-04 | 1.5 | 1.14E-01 |
| GOTERM_MF_DIRECT | ***sr-e*** | Organic cation transmembrane transporter activity | 9 | 1.16E-08 | 18.0 | 3.17E-06 |
| GOTERM_BP_DIRECT |  | **Cellular response to heat** | 6 | 2.07E-05 | 16.3 | 1.52E-02 |
| INTERPRO |  | **Stress-inducible humoral factor Turandot** | 5 | 2.94E-05 | 23.8 | 1.29E-02 |
| GOTERM_BP_DIRECT |  | **Response to oxidative stress** | 11 | 8.01E-05 | 4.9 | 2.92E-02 |
| GOTERM_MF_DIRECT |  | Carboxypeptidase activity | 5 | 2.72E-04 | 14.6 | 3.66E-02 |
| GOTERM_BP_DIRECT |  | **Transmembrane transport** | 19 | 2.76E-04 | 2.7 | 6.59E-02 |
| GOTERM_BP_DIRECT |  | **Response to UV** | 5 | 3.86E-04 | 13.6 | 6.90E-02 |
| GOTERM_MF_DIRECT |  | **Photoreceptor activity** | 4 | 9.00E-04 | 19.0 | 7.89E-02 |

Relevant to seasonal adaptations and desiccation tolerance categories are marked by bold type
