## Supplementary material for "Seasonal changes in recombination rate, crossover interference, and their response to desiccation stress in a natural population of *Drosophila melanogaster* from India": Table S2

**Table S2. Spearman correlations between the desiccation-induced changes in recombination rate and desiccation tolerance**

| Interval | Absolute changes in recombination rate | | | | Relative changes in recombination rate | | | |
| --- | --- | --- | --- | --- | --- | --- | --- | --- |
|  | Desiccation tolerance  of the hybrid | | Desiccation tolerance  of the parental line | | Desiccation tolerance  of the hybrid | | Desiccation tolerance  of the parental line | |
|  | ρ | *p* | ρ | *p* | ρ | *p* | ρ | *p* |
| Chromosome X | | | | | | | | |
| *y-cv* | -0.659 | 0.010 | -0.640 | 0.014 | -0.665 | 9.4·10^-3^ | -0.631 | 0.016 |
| *cv-v* | -0.467 | 0.092 | -0.455 | 0.102 | -0.485 | 0.079 | -0.486 | 0.078 |
| *v-f* | -0.407 | 0.148 | -0.336 | 0.240 | -0.407 | 0.148 | -0.336 | 0.240 |
| Chromosome 2 | | | | | | | | |
| *al-dp* | -0.476 | 0.086 | -0.455 | 0.102 | -0.480 | 0.082 | -0.468 | 0.091 |
| *dp-b* | 0.344 | 0.229 | 0.204 | 0.483 | 0.348 | 0.223 | 0.200 | 0.493 |
| *b-pr* | 0.163 | 0.578 | 0.200 | 0.493 | 0.088 | 0.765 | 0.125 | 0.670 |
| *pr-c* | -0.751 | 2.0·10^-3^ | -0.609 | 0.021 | -0.751 | 2.0·10^-3^ | -0.609 | 0.021 |
| *c-px* | -0.337 | 0.239 | -0.288 | 0.318 | -0.476 | 0.086 | -0.393 | 0.164 |
| *px-sp* | 0.634 | 0.015 | 0.692 | 6.1·10^-3^ | 0.676 | 7.9·10^-3^ | 0.692 | 6.1·10^-3^ |
| Chromosome 3 | | | | | | | | |
| *ru-h* | -0.692 | 6.1·10^-3^ | -0.626 | 0.017 | -0.710 | 4.5·10^-3^ | -0.648 | 0.012 |
| *h-th* | -0.341 | 0.233 | -0.380 | 0.180 | -0.411 | 0.144 | -0.451 | 0.106 |
| *th-sr* | -0.182 | 0.533 | -0.279 | 0.334 | -0.213 | 0.464 | -0.301 | 0.296 |
| *sr-e* | 0.174 | 0.553 | 0.222 | 0.446 | 0.169 | 0.563 | 0.213 | 0.464 |
