## Supplementary material for "Seasonal changes in recombination rate, crossover interference, and their response to desiccation stress in a natural population of *Drosophila melanogaster* from India": Table S3

**Table S3. Spearman correlations between the desiccation-induced changes in crossover interference and desiccation tolerance**

| Pairs of  intervals | Absolute changes in crossover interference | | | | Relative changes in crossover interference | | | |
| --- | --- | --- | --- | --- | --- | --- | --- | --- |
|  | Desiccation tolerance  of the hybrid | | Desiccation tolerance  of the parental line | | Desiccation tolerance  of the hybrid | | Desiccation tolerance  of the parental line | |
|  | ρ | *p* | ρ | *p* | ρ | *p* | ρ | *p* |
| Chromosome X | | | | | | | | |
| *y–cv–v* | -0.170 | 0.562 | -0.002 | 0.994 | -0.229 | 0.431 | -0.029 | 0.923 |
| *y–v–f* | -0.659 | 0.010 | -0.521 | 0.056 | -0.676 | 7.9·10^-3^ | -0.495 | 0.072 |
| *cv–v–f* | -0.639 | 0.014 | -0.543 | 0.045 | -0.643 | 0.013 | -0.464 | 0.095 |
| Chromosome 2 | | | | | | | | |
| *al–dp–b* | -0.344 | 0.229 | -0.437 | 0.118 | -0.119 | 0.685 | -0.284 | 0.326 |
| *al–dp–pr* | -0.515 | 0.059 | -0.569 | 0.034 | -0.498 | 0.070 | -0.587 | 0.027 |
| *al–b–c* | -0.328 | 0.252 | -0.402 | 0.154 | -0.337 | 0.239 | -0.393 | 0.164 |
| *dp–b–c* | -0.073 | 0.805 | -0.143 | 0.626 | -0.112 | 0.702 | -0.204 | 0.483 |
| *dp–pr–c* | -0.037 | 0.899 | -0.138 | 0.637 | -0.084 | 0.776 | -0.182 | 0.533 |
| *b–c–px* | 0.247 | 0.395 | 0.125 | 0.670 | 0.207 | 0.478 | 0.090 | 0.759 |
| *b–c–sp* | 0.253 | 0.382 | 0.130 | 0.659 | 0.218 | 0.454 | 0.090 | 0.759 |
| *pr–c–px* | 0.293 | 0.309 | 0.213 | 0.464 | 0.337 | 0.239 | 0.235 | 0.418 |
| *pr–c–sp* | 0.322 | 0.262 | 0.169 | 0.563 | 0.313 | 0.276 | 0.156 | 0.594 |
| Chromosome 3 | | | | | | | | |
| *ru–h–th* | 0.393 | 0.164 | 0.389 | 0.169 | 0.420 | 0.135 | 0.420 | 0.135 |
| *h–th–sr* | 0.336 | 0.240 | 0.279 | 0.334 | 0.380 | 0.180 | 0.327 | 0.253 |
| *h–th–e* | 0.508 | 0.064 | 0.547 | 0.043 | 0.490 | 0.075 | 0.556 | 0.039 |
| *th–sr–e* | -0.165 | 0.573 | -0.138 | 0.637 | -0.169 | 0.563 | -0.134 | 0.648 |
